# Soil pH influences the composition of bacteriophage communities infecting individual hosts

**DOI:** 10.1101/2024.10.04.616616

**Authors:** Sungeun Lee, Graeme W. Nicol, Christina Hazard

## Abstract

Bacteriophages (phages) can infect a range of hosts in highly diverse soil bacterial communities. However, selection of host communities across ecological gradients and co-evolutionary processes may influence both the distribution of phages and the susceptibility of individual hosts through virus interactions and local adaptation within distinct ecological niches. Metagenomic-based analyses have revealed that soil pH selects for distinct populations and community structures for both phage and hosts at local and global scales. However, whether contrasting soil pH represents a selective barrier for phages capable of infecting an individual host is unknown. To examine the influence of pH on individual host-virus interactions, two closely related *Bacillus* strains were isolated and characterized from pH 7.5 soil associated with a long-term contiguous pH gradient (pH 4.5 to 7.5). Phages infecting each strain were subsequently enriched from soils across the pH gradient (pH 4.5, 5.5, 6.5 and 7.5), enumerated using a plaque assay, and characterized via metagenomic analysis. Phages infecting each strain were cultivated from all soils but their community composition and abundance varied with pH. Phage populations infecting each of the two strains were distinct despite the close relatedness of the two hosts, indicating relatively narrow host ranges for each virus. These results suggest that while phage community structures vary substantially across an ecological gradient, soil pH alone does not represent a barrier for the distribution of phages capable of infecting an individual host.

## Introduction

The soil virosphere is highly diverse, active and spatiotemporally dynamic with implications for microbial communities and ecosystem function (Kuzyakov and Mason-Jones, 2018; Pratama and van Elsas, 2018; Roux and Emerson, 2022). Viruses modulate microbial abundance and metabolic potential through infection and cell lysis with these actions constituting a major determinant of microbial evolution and ecology (Koskella and Brockhurst, 2014; Williamson et al., 2017; Pratama et al., 2021). Susceptibility of microbial hosts to viral infection depends on physical encounter, but also viral attachment, counter-defence and resistance strategies and local adaptation, which are tightly controlled through coevolutionary processes (de Jonge et al., 2019; Crespo-Bellido and Duffy, 2023).

Infectivity of soil phages have been shown to be tightly coupled to bacterial hosts at the cm scale in field studies (Vos et al., 2009). In laboratory microcosms, rapid changes in soil virus richness and composition has been demonstrated, with viral predation of the dominant bacterial hosts suggesting ‘cull the winner’ dynamics (Santos-Medellín et al., 2023). High estimates of viral driven microbial death in soil have also been predicted (Nicolas et al., 2023). At the field scale, spatial distance versus abiotic factors have been observed to have a stronger impact on soil virus community structure (Santos-Medellín et al., 2022), though large variation in soil pH has been shown to influence virus community structure at the local and global scale as typically observed for prokaryotes (Bartram et al., 2014; Adriaenssens et al., 2017; Bahram et al., 2018; Lee et al., 2022). Correspondingly, similar abundance patterns between hosts and viruses has repeatedly been found based in metagenomic analyses (Srinivasiah et al., 2015; Coclet et al., 2023), suggesting linked host and virus population dynamics. While changes in virus community structure are observed with ecological gradients in metagenomic studies, these typically do not identify how infection of individual hosts varies or whether changes in virus community structure reflects changes in host susceptibility to infection.

In a previous metagenome-based study (Lee et al., 2022), we observed that that pH 4.5 and 7.5 soils from contrasting ends of a contiguous pH gradient contained largely separate virus communities with 524 and 1,361 viral operational taxonomic units (vOTUs) detected exclusively in either soil, respectively, and represented 99% of all vOTUs. Using the same soil samples, here we investigated whether these contrasting community structures indicates that soil pH represents a barrier to the distribution of viruses infecting an individual host. Native bacterial strains were isolated and infecting lytic phages present in soils from across the pH gradient enumerated using plaque assays and the identity and relative abundance of phage determined through metagenomic sequencing approaches.

## Material and Methods

### Soil sampling

Soil was collected from the long-term pH gradient (pH 4.5 to 7.5 at 0.5 intervals) at Craibstone Research Station, SRUC, Aberdeen, Scotland (57.185, -2.212) in January 2019 (see Lee et al. (2022) for detailed description). Individual soil samples were collected in triplicate at 10 cm depth at 1 m intervals within pH 4.5, 5.5, 6.5 and 7.5 subplots. Soil samples were sieved to 2 mm and stored at 4°C.

### Isolation and selection of host strains for plaque assay

Bacterial strains were isolated on 1/10 tryptone soy agar (TSA) plates from pH 7.5 soil after adding 1 g sieved soil to 9 ml of phosphate-buffered saline, manually shaking for 1 min, and spread-plating 100 µl of serial dilutions (10^-1^, 10^-2^ and 10^-3^) before incubating at 25°C for 3 days. A total of 60 colonies were picked and streaked onto fresh TSA plates to confirm isolation. All 60 isolated strains were then screened for the ability to form confluent growth and demonstrate susceptibility to lytic infection from soil using plaque assays (described below). Only nine of the 60 strains met both criteria. Bacterial colonies from each of these nine strains were used for direct PCR amplification of near full-length 16S rRNA genes with primers pA and pH (Edward et al., 1989). Each 50 μl PCR reaction contained 10x buffer, 0.4 µM of each primer, 2 mM dNTP, 1 unit Taq DNA polymerase (Thermo Fisher) and PCR grade water, and used the following thermal cycling program: 1 cycle of 10 min at 95°C; 30 cycles of 30 s at 95°C, 30 s at 55°C and 30 s at 72°C; 1 cycle of 5 min at 72°C. PCR products were Sanger sequenced (Eurofins Genomics) before comparison to sequences in NCBI GenBank by BLASTn searches. All strains had >98% identity to known *Bacillus* strains with two (S4 and C30) selected for subsequent experiments.

### *Genome sequencing of* Bacillus *S4 and C30 strains*

Strains were grown in 20 ml 1/10 tryptic soy broth (TSB) liquid medium at 29°C overnight. Cells were pelleted by centrifugation at 6,000 x *g* for 10 min and supernatant removed before resuspension in 500 µl of Tris-EDTA (TE) buffer. Cells were lysed by incubating with 20 µl of lysozyme (100 mg ml^-1^) (Sigma-Aldrich) at 37°C for 30 min. The lysate was further incubated at 56°C for 1 h after the addition of 40 µl of 10% sodium dodecyl sulfate (SDS) and 8 µl of proteinase K solution (100 mg ml^-1^) (Promega), followed by CTAB extraction, chloroform extraction and PEG precipitation as previously described (Nicol and Prosser, 2011). DNA quality was assessed by agarose gel electrophoresis and quantified using a Qubit dsDNA HS Assay kit (Invitrogen).

DNA was diluted to a concentration of 0.25 ng µl^-1^ and libraries prepared with 1 ng DNA using the Nextera XT Library Prep Kit and sequenced on a MiSeq platform (Illumina) with 250 bp paired-end reads. Quality filtering of raw sequences (S4, 1.46 M reads; C30, 1.27 M reads) was performed using metaWRAP read_qc module (Uritskiy et al., 2018). Contig assembly was done using metaSPAdes (Nurk et al., 2017) and binning with metaWRAP bin_refinement module. Taxonomic classification of genomes was carried out using GTDB-Tk v0.3.2 (Chaumeil et al., 2019) and genome completeness assessed with CheckM (Parks et al., 2015). Phylogenomic analysis was performed using GtoTree (Lee, 2019). Prophage detection was predicted using PhageBoost (Sirén et al., 2020) and viral quality verified using CheckV (Nayfach et al., 2021). CRISPR arrays were searched for using the CRT tool (Bland et al., 2007) and DefenceFinder (Tesson et al., 2022) to detect antiviral systems.

To investigate the distribution of closely-related *Bacillus* strains with identical 16S rRNA gene sequences (V4 region only) to S4 and C30 across the pH gradient, 16S rRNA gene amplicon data previously derived from the same soil pH gradient was analysed (Bartram et al., 2014) SRA accession no. SRX352269, run SRR988432)). Identical sequence reads from each soil pH were mapped to the 16S rRNA gene of *Bacillus* S4 and C30 strains using BBMap (Bushnell, 2014).

### Plaque assays

An overview of the enrichment, cultivation, enumeration and metagenome sequencing of phage is provided in Fig. S1. Phage populations infecting *Bacillus* S4 and C30 were enriched from triplicate field soil samples of each pH (pH 4.5, 5.5, 6.5 and 7.5) by combining 0.2 g dry-weight equivalent soil and 100 µl of overnight grown culture in 1 ml of 1/10 TSB and incubated at 29°C for 16 h. The medium was then filtered through 0.2 µm syringe filters and the filtrate serially diluted to 10^-1^, 10^-2^ and 10^-3^ (n = 48) with phage buffer (100 mM NaCl, 8 mM MgSO_4_×7H_2_O, 50 mM Tri-HCl, pH 7.5 and 0.02 µm filtered). The influence of soil pH on the growth of each strain was assessed by measuring changes in optical density (OD) with no significant differences observed between the different soils (Fig. S2).

Plaque assays were performed using virus enrichments from each pH soil applied to *Bacillus* S4 and C30 cultures. Confluent growth on agar plates was obtained by adding 500 µl late exponential phase culture combined with 4.5 ml 1/10 TSB soft agar (3 g agar l^-1^) with either 10 µl of 10^0^, 10^-1^, 10^-2^ or 10^-3^ viral enrichment or 1/10 TSB medium as a negative control. Samples were incubated for 5 min at room temperature and poured on top of 1/10 TSA plates and dried, sealed with parafilm, and incubated at 25°C for two days. A total of 60 plates were produced for each strain (three replicates x four pH soils x four virus enrichment dilutions and negative control). The 10^-3^ viral enrichment plates (<200 PFUs) were used to determine the number of plaque forming units (PFU) with 44 x10^3^ to 303 x10^6^ PFU ml^-1^ produced for all treatments.

To recover lytic phage populations from each soil pH, total phages were washed from triplicate 10^-3^ viral enrichment plates (12 per strain). Specifically, 2 ml of phage buffer was added onto the surface of each plate, agitated by gentle shaking at 50 rpm at 25°C for 2 h, and removed before passed through 0.2 µm to remove *Bacillus* cells. The phage wash was used for transmission electron microscopy (TEM) and metagenomic sequencing.

### Transmission electron microscopy of phages

Phage populations obtained from pH 4.5 and 7.5 plaque assays were prepared for TEM by placing a sample of plate wash onto 200 mesh-size TEM nickel grids coated in formvar-C (EMS, Hatfield, USA) and leaving to absorb for 2 min. Grids were then stained with 2% phosphotungstic acid for 2 min before viewing with a JEM-1400 120 kV transmission electron microscope (Jeol) equipped with a Orius 600 camera (Gatan) and Digital Micrograph Software (Gatan).

### Metagenomic characterization of phage populations

Phage DNA was extracted from the 12 individual filtered plate washes. Samples were treated with DNase I to remove free DNA (RQ1 DNase; Promega) and lysed by adding 40 µl of 10% SDS and 4 µl of of proteinase K (100 ug ml^-1^). Lysate was incubated at 37°C for 1 h followed by CTAB extraction, chloroform separation and PEG precipitation as previously described (Nicol and Prosser, 2011). DNA quality assessment, library preparation, and MiSeq sequencing was as described above for *Bacillus* strain genome sequencing. Long-read sequencing was also performed (Overholt et al., 2020) using an equimolar pool of DNA from each sample on a MinION sequencer (Oxford Nanopore Technologies). The 1D Genomic DNA by Ligation Kit (SQK-LSK109) was used for library preparation following manufacturers’ protocol with minor modifications. The ‘end-prep’ step was performed with 50 µl of DNA sample, 0.5 µl of DNA CS, 7 µl of Ultra II End-prep reaction buffer, and 3 µl of Ultra II End-prep enzyme mix. Samples were incubated at 20°C followed by 65°C for 5 min using a thermal cycler. The product was purified with 1X AMPure XP beads (Beckman Coulter) and eluted in 31 µl of nuclease-free water. An adapter was ligated by mixing 30 µl of DNA, 50 µl of NEB Blunt/TA Ligase Master Mix (New England Biolabs), 5 µl of adapter mix (AMX) and 15 µl of nuclease-free water and incubated for 15 min at room temperature. Purification and loading of the libraries were performed according to manufacturers’ protocols and sequenced using the MinKNOW FLO-MNI106 workflow.

Viromes were checked for host contamination by mapping reads to the assembled host genome using Salmon mapper v0.9.1 (Patro et al., 2017). Quality filtering of raw sequences was performed using the metaWRAP read_qc module (Uritskiy et al., 2018). Quality-controlled MiSeq reads (S4 phages, 14 M reads; C30 phages, 20 M reads) and Nanopore reads (S4 phages, 2,138 reads; C30 phages, 349 reads) were used to produce long-read contigs with Unicycler (Wick et al., 2017). Contigs >10 kb were used in downstream analyses. Phage contigs were compared to reference virus genomes from Virus-Host DB (Mihara et al., 2016) using VIPTree (Nishimura et al., 2017). Taxonomic classification of phage contigs was performed using geNomand (Camargo et al., 2023), gene prediction with Prodigal (Hyatt et al., 2010) and homology searches using InterProScan 5 (E value < 10-5) (Jones et al., 2014). Comparison of spacer sequences in CRISPR arrays in host genomes to phage reads and contigs was performed using the locate function of Seqkit (Shen et al., 2016) and BLASTn with blastn-short option. Normalized relative abundance was calculated using Quant_bins module in Salmon v0.9.1 (Patro et al., 2017; Uritskiy et al., 2018).

## Results

### Isolation and characterization of host strains

Two strains (*Bacillus* S4 and C30) were chosen that possessed the two desired criteria of being able to form confluent growth and demonstrating susceptibility to lytic viral infection in plaque assays. Taxonomic classification of the high-quality genomes of S4 and C30 (95% and 92% estimated completeness, respectively; ≤0.9% contamination) had 97.2% and 95.0% average nucleotide identity (ANI) to *Bacillus cereus* and 95.1% and 97.9% ANI to *Bacillus mycoides*, respectively. Phylogenomic analysis demonstrated that both strains represented the same *Bacillus* species sharing 96% ANI (Rodriguez-R and Konstantinidis, 2016) and possessed only 1 bp difference in full-length 16S rRNA gene sequences which were identical over the V4 region analysed in amplicon-based 16S rRNA gene surveys. The presence of *Bacillus* strains possessing an identical partial (V4) 16S rRNA sequence was confirmed in pH 4.5, 5.5, 6.5 and 7.5 soils using 16S rRNA gene amplicon sequencing with the total proportion of ASVs increasing with pH and ranging from 0.035 - 0.111% (Fig. S3). Both strains had evidence of antiviral systems including CRISPR-Cas and restriction modification (RM) system Type IV, and abortive infection (Abi) systems AbiJ and AbiQ in *Bacillus* S4.

### Host infectivity by lytic phage in pH 4.5 to 7.5 soils

Both *Bacillus* hosts were infected by viruses in soils from across the entire pH gradient (Fig. 1). A significant trend of decreasing PFUs with increasing pH was found for both strains (r2 >0.58, *p*-value <0.003) with the highest number of PFU recovered from pH 4.5 soil (1.1 x 10^8^ (s.e. 0.3) and 0.7 x 10^8^ (s.e. 0.1) PFU ml^-1^ for C30 and S4, respectively) and were lowest in pH 7.5 soil (1.1 x 10^8^ (s.e. 0.3) and 0.7 x 10^8^ (s.e. 0.1) PFU ml^-1^ for C30 and S4, respectively). The number of PFUs were significantly higher for C30 compared to S4 at all pH (4, 5, 6 and 89-fold higher at pH 4.5, 5.5, 6.5 and 7.5, respectively).

**Fig. 1:**
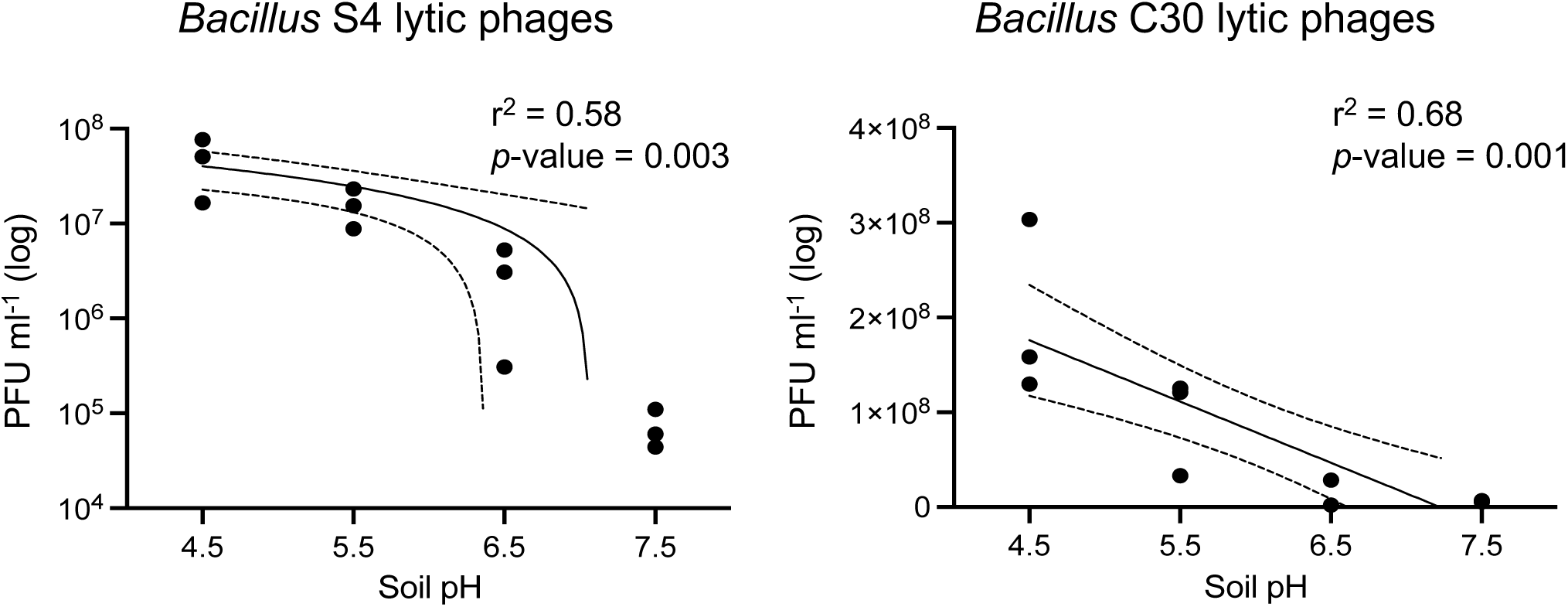
Abundance of lytic phage infecting *Bacillus* S4 and C30 enriched from across a pH 4.5 to 7.5 soil pH gradient as determined from plaque assays. Regression line, 95% confidence interval of the regression line, and associated statistics are given.

### Characterization of lytic phage populations infecting host strains

Viruses were washed off agar plates with plaques for metagenomic sequencing of each replicate. TEM confirmed phage-like particles in the plaque assay washes (Fig. S4). From all plates, eight full length virus genomes were recovered with three infecting S4 and five infecting C30 (Fig. 2). All were closely related to previously characterized *Bacillus* phages within the *Caudoviricetes* and were not derived from integrated prophages. No phage genome sequence matched spacers in CRISPR arrays with some regions only sharing partial identity over a reduced portion (i.e. 11-29 bp of genomic regions matching spacers 39-40 bp spacers) (Table S1). No individual phage infected both strains (Fig. 2). For both strains, one individual phage represented 100% of PFUs from pH 4.5 soils (S4_phage_1) or pH 4.5 and 5.5 soils (C30_phage_1), respectively, with both belonging to the family *Herelleviridae* (Fig. 3). While these two viruses were present in soils at all pH, the relative abundance of reads mapped to viruses from other families increased with increasing pH, with S4_phage_2 and C30_phage_2 representing 56% (s.e. 10%) and 28% (s.e. 18%) at pH 7.5.

**Fig. 2:**
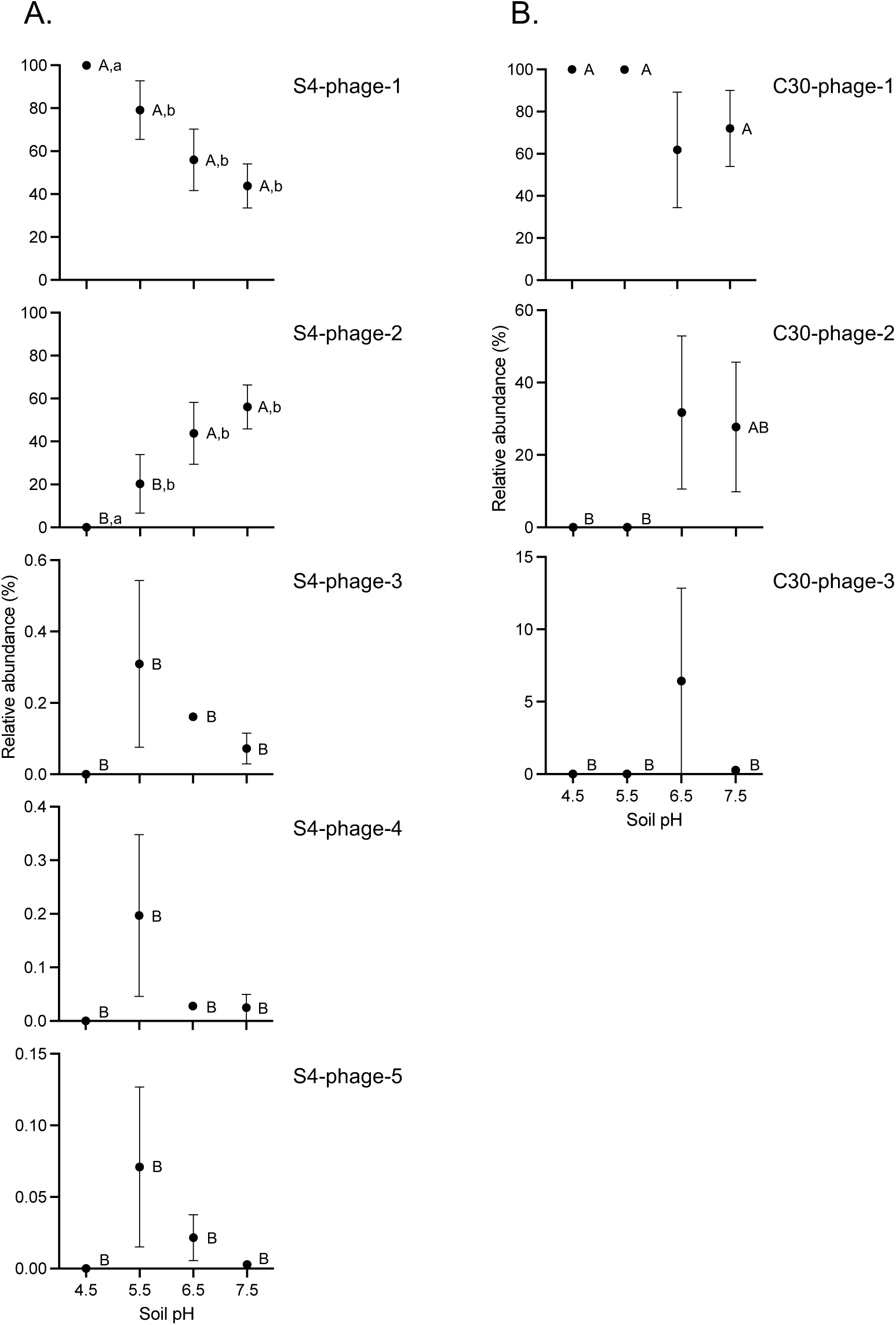
Relative abundance of enriched lytic phages infecting A) *Bacillus* S4 and B) *Bacillus* C30 strains inferred from the relative abundance of reads mapped to phage genomes after correcting for genome length. For each individual soil pH, significantly different relative abundances between different phage infecting one individual strain are denoted with different uppercase letters. For each individual phage, significantly different relative abundances associated with changing pH are denoted with lowercase letters (Tukey’s test *p*-value <0.05).

**Fig. 3:**
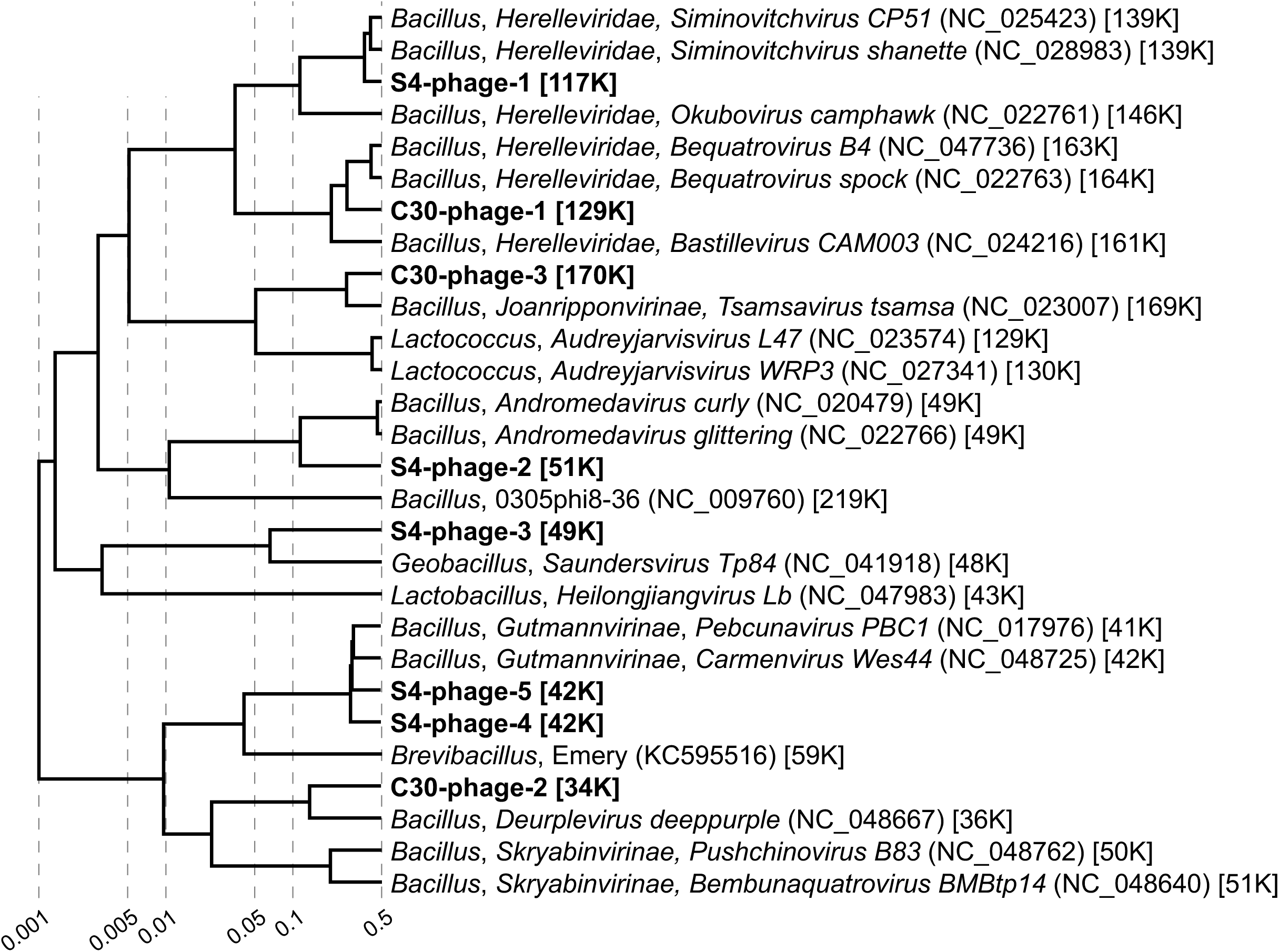
Proteomic tree using whole genome sequences showing relationships of S4 and C30-infecting viruses with reference phages infecting other *Bacillus* or class *Bacilli* strains. Names denote genus of host, family or sub-family lineage of virus (if classified by the International Committee on Taxonomy of Viruses), and virus name, with details in parentheses and brackets denoting NCBI accession number and genome length, respectively. Values at dotted lines represent a distance metric based on normalized tBLASTx scores plotted on a log scale.

## Discussion

The aim of this study was to determine if viruses capable of infecting an individual host bacterium are restricted to a particular pH across a contiguous soil pH gradient and if pH represents a barrier to their distribution. The two *Bacillus* host strains used in this study originated from pH 7.5 soil and were isolated on neutral pH medium. The increase in the abundance of viruses infecting both strains with decreasing soil pH is perhaps surprising considering that previous work has demonstrated that host-virus interactions and susceptibility to infection is higher when host and virus are co-localized over relatively short (∼cm) scales (Vos et al., 2009). This may indicate that co-localization of viruses and hosts in pH 7.5 soil results in adaptation of host resistance to native virus populations, and viruses in acidic soils, spatially isolated through a selective barrier of pH-associated niche separation, have greater infectivity through reduced previous interactions that would lead to increased resistance. Alternatively, it is possible that the lower abundance of PFUs from pH 7.5 soil is due to co-localization resulting in faster rates of infection and lysis. During the enrichment stage, more efficient rates of infection could result in a lower titre of virions that are subsequently enumerated in plaque assays, with slower rates of interaction, infection or reproduction associated with non-co-localized (i.e. low pH soil) viruses resulting in a higher net production of virions.

The use of a pre-incubation step to enrich for phage prior to application on lawns of the same strain was required to obtain sufficient PFUs for reproducible enumeration. However, this also removed our ability to directly enumerate the number of viable phages in the soil. The reproducible and significant trends of changes in abundance of enriched infective phages across the pH gradient for both strains indicates that this approach can reflect real differences in virus distribution and abundance. The addition of soil during the enrichment stage did impact *Bacillus* growth but culture OD was not significantly different for soils of contrasting pH and was therefore unlikely a source of differences in enumerated viruses.

The two *Bacillus* strains used in this study do not represent different species based on ANI. Phages infecting each strain were distinct and therefore suggests host specificity was restricted to the strain/sub-species level with a narrow host range. Comparatively, previous studies of *Bacillus* viruses isolated from soil and wastewater have shown host ranges at the genus, species and strain level based on plaque assays, spot tests and/or culture OD, and using host strains isolated from different environmental samples as that of the virus (El-Arabi et al., 2013; Krasowska et al., 2015; Sauder et al., 2016; Willms et al., 2017; Li et al., 2020).

Both *Bacillus* strains had common antiviral systems. Prokaryotes encode on average five antiviral systems with RM (83%) and CRISPR-Cas (39%) being the most abundant (Tesson et al., 2022). The phages that infected the hosts did not match spacers in the CRISPR arrays of the hosts, but did match at a much lower coverage, likely reflecting CRISPR ‘dark matter’ versus point mutations allowing for viral escape (Shmakov et al., 2020; Rubio et al., 2023). One of the host strains also had two different abortive infection systems, perhaps contributing to the different viral populations that infected the two host strains.

Due to the vast diversity of hosts and viruses in soils it is challenging to study individual virus-host dynamics using metagenomic analyses alone (Roux and Emerson, 2022). This *in vivo* plaque assay approach enabled identifying the distribution and abundance of viruses infecting focal native populations across a soil pH gradient. In the gradient soil samples used in this study, virus community structures were previously found to be strongly influenced by soil pH with almost no overlap in the detection of vOTUs identified in pH 4.5 and 7.5 soils. While one S4 and one C30 infecting phage were recovered from all soils, the results presented here are congruent with these previous findings with different virus communities reproducibly enriched in soils of contrasting pH. They also demonstrate that separate and largely unique metagenome-derived virus community structures between different soils does not reflect separation of the infectivity potential of a virus community towards an individual strain.

## Data availability

Metagenome draft assemblies are accessible in NCBI’s GenBank under BioProject accession nos. PRJNA1136417 (*Bacillus* S4 and C30 strains) and PQ069201 – PQ069208 (phage genomes).

## Acknowledgments

The authors would like to thank Lucas P. P. Braga and Laurent Philippot for practical guidance with the plaque assay approach and acknowledge use of the Newton high-performance computing cluster at PMCS2I, Ećole Centrale de Lyon. This work was funded by the AXA Research Fund.

## Supplementary Information

### 1. Supplementary Table

**Table S1:**
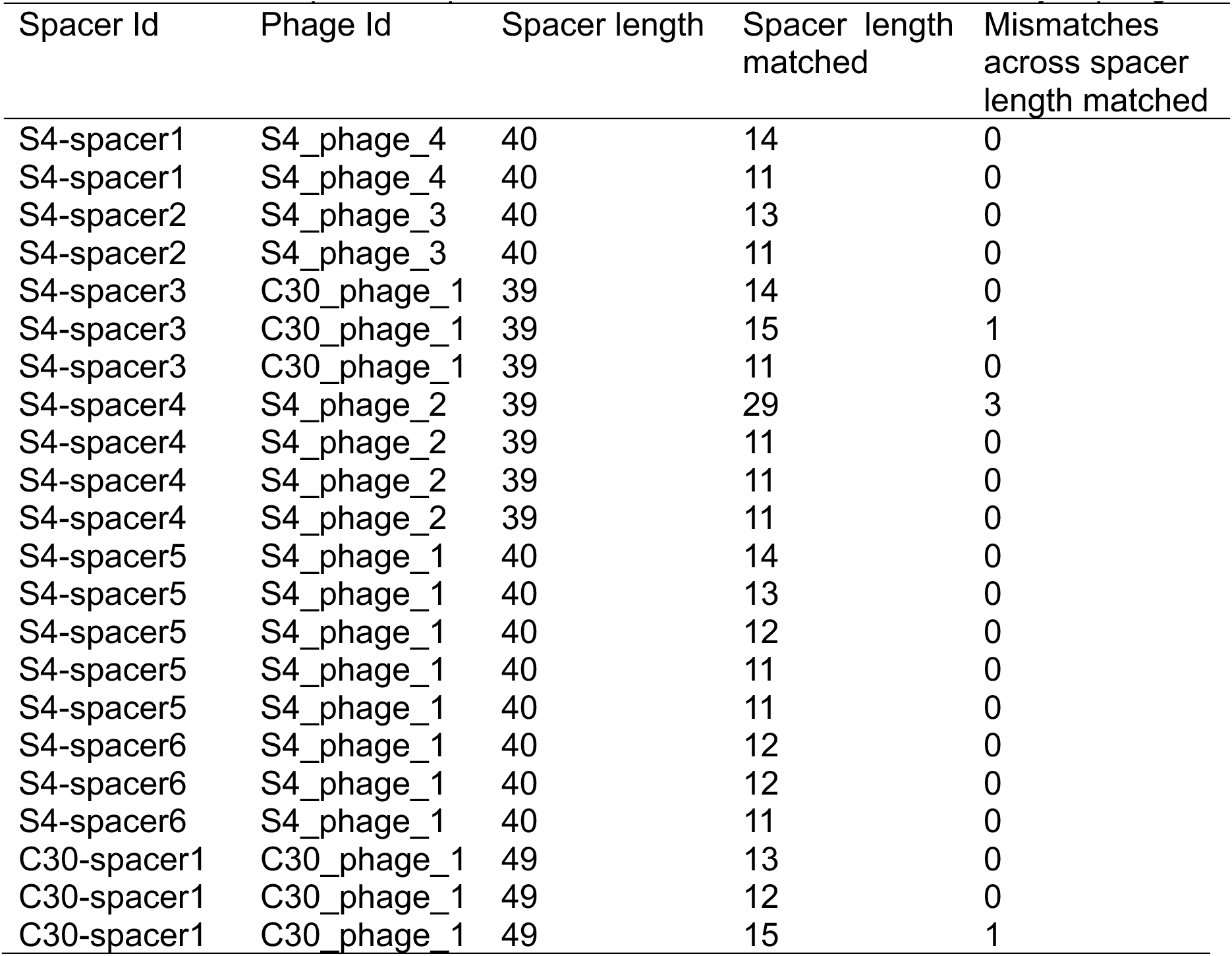
CRISPR spacer sequence matches to *Bacillus* S4 and C30 lytic phages.

| Spacer Id | Phage Id | Spacer length | Spacer length<br>matched | Mismatches<br>across spacer<br>length matched |
| --- | --- | --- | --- | --- |
| S4-spacer1 | S4_phage_4 | 40 | 14 | 0 |
| S4-spacer1 | S4_phage_4 | 40 | 11 | 0 |
| S4-spacer2 | S4_phage_3 | 40 | 13 | 0 |
| S4-spacer2 | S4_phage_3 | 40 | 11 | 0 |
| S4-spacer3 | C30_phage_1 | 39 | 14 | 0 |
| S4-spacer3 | C30_phage_1 | 39 | 15 | 1 |
| S4-spacer3 | C30_phage_1 | 39 | 11 | 0 |
| S4-spacer4 | S4_phage_2 | 39 | 29 | 3 |
| S4-spacer4 | S4_phage_2 | 39 | 11 | 0 |
| S4-spacer4 | S4_phage_2 | 39 | 11 | 0 |
| S4-spacer4 | S4_phage_2 | 39 | 11 | 0 |
| S4-spacer5 | S4_phage_1 | 40 | 14 | 0 |
| S4-spacer5 | S4_phage_1 | 40 | 13 | 0 |
| S4-spacer5 | S4_phage_1 | 40 | 12 | 0 |
| S4-spacer5 | S4_phage_1 | 40 | 11 | 0 |
| S4-spacer5 | S4_phage_1 | 40 | 11 | 0 |
| S4-spacer6 | S4_phage_1 | 40 | 12 | 0 |
| S4-spacer6 | S4_phage_1 | 40 | 12 | 0 |
| S4-spacer6 | S4_phage_1 | 40 | 11 | 0 |
| C30-spacer1 | C30_phage_1 | 49 | 13 | 0 |
| C30-spacer1 | C30_phage_1 | 49 | 12 | 0 |
| C30-spacer1 | C30_phage_1 | 49 | 15 | 1 |

### 2. Supplementary Figures

**Fig. S1:**
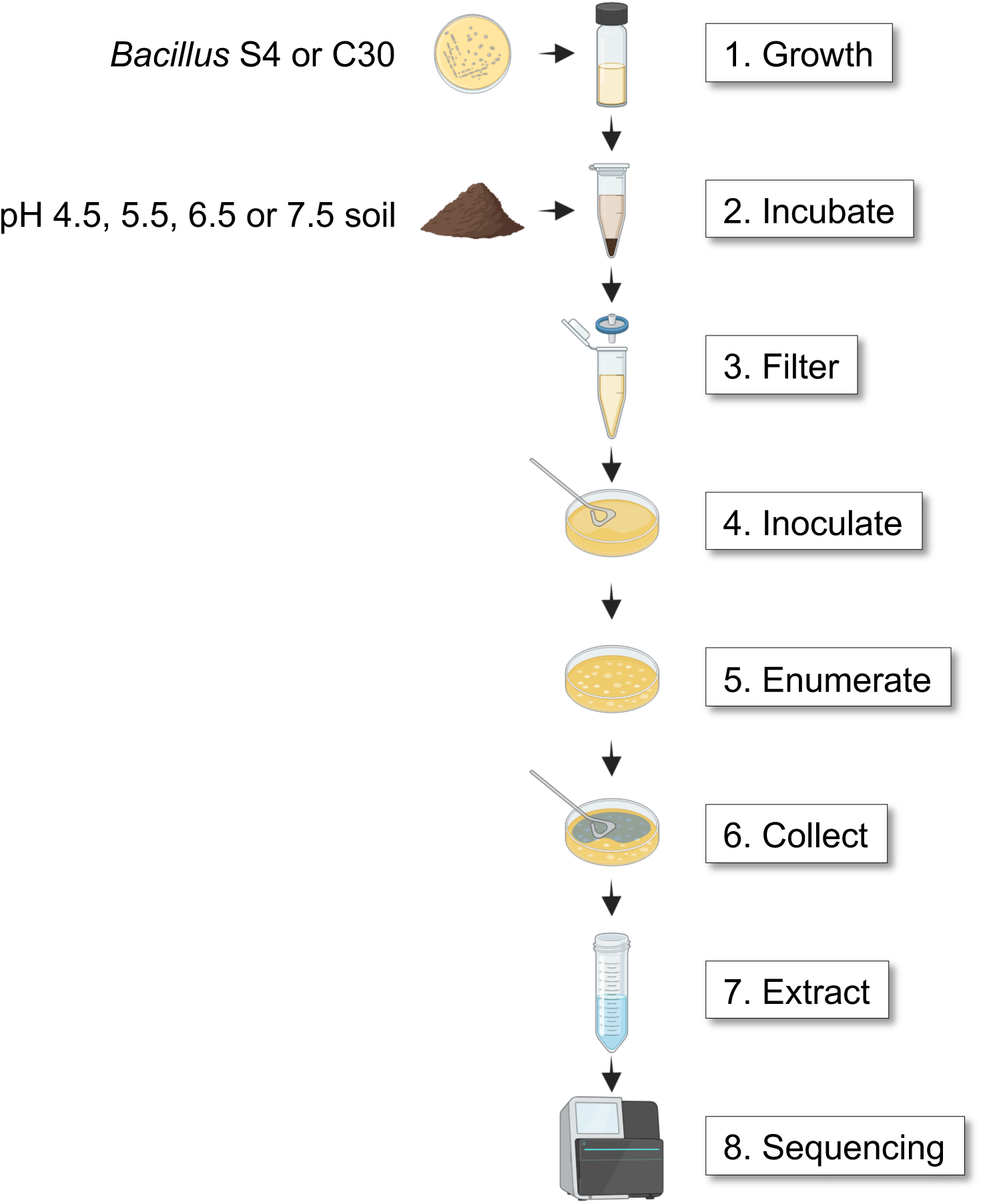
Overview of analysis of lytic phages from soil infecting *Bacillus* S4 and C30. 1) Overnight cultures of each strain were prepared from an isolated colony. 2) 0.2 g soil (dry-weight equivalent) and 100 µl of overnight grown culture added to 1 ml of 1/10 TSB and incubated at 29°C for 16 h. 3) Medium filtered through 0.2 µm filter and serially diluted. 4) Phage-enriched filtrates added to soft agar plates with *Bacillus* S4 or C30. 5) PFUs are counted at an appropriate dilution (<200 PFUs per plate). 6) Phage particles are washed and recovered from plates. 7) DNA extracted libraries prepared for sequencing. 8) Phage metagenomes sequenced using MiSeq (Illumina) or MinION (Oxford Nanopore) sequencing.

**Fig. S2:**
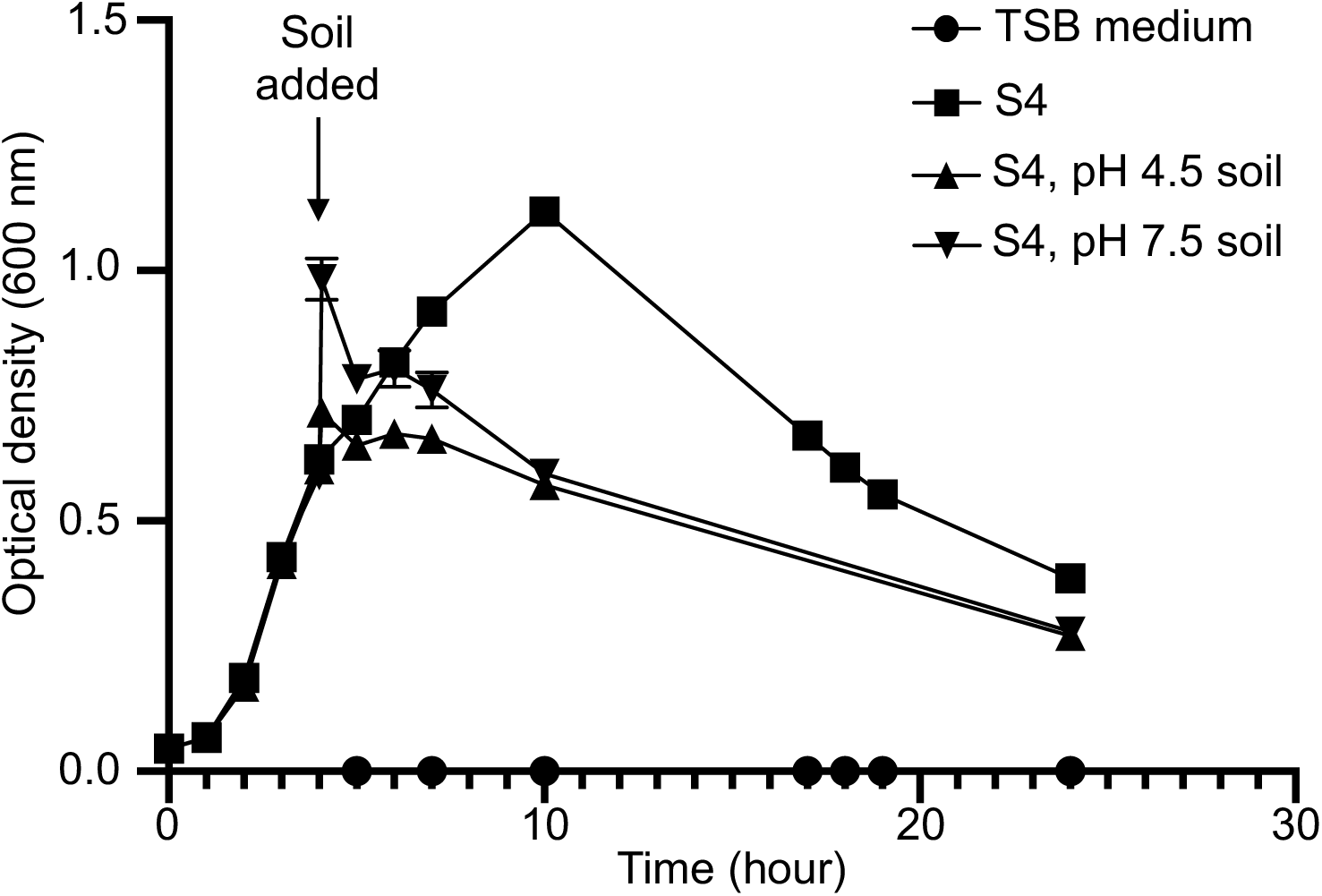
Growth and optical density of *Bacillus* S4 cultures after the addition of pH 4.5 or 7.5 soil (5 g (dry weight equivalent) in 25 ml culture) during exponential growth in tryptone soya broth medium. Error bars represent standard error of the mean of triplicate cultures. Optical density was measured using a NanoPhotometer (Implen).

**Fig. S3:**
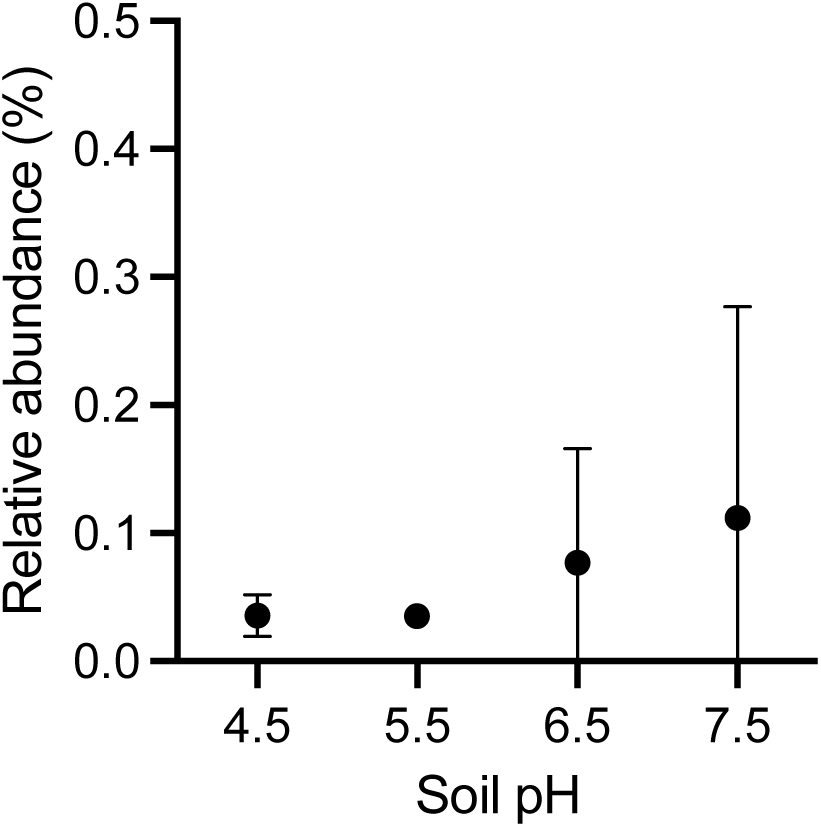
Relative abundance of *Bacillus* strains possessing an identical partial (V4) 16S rRNA sequence to that of *Bacillus* S4 and C30 strains across the soil pH gradient. Error bars represent standard error of the mean of triplicate samples.

**Fig. S4:**
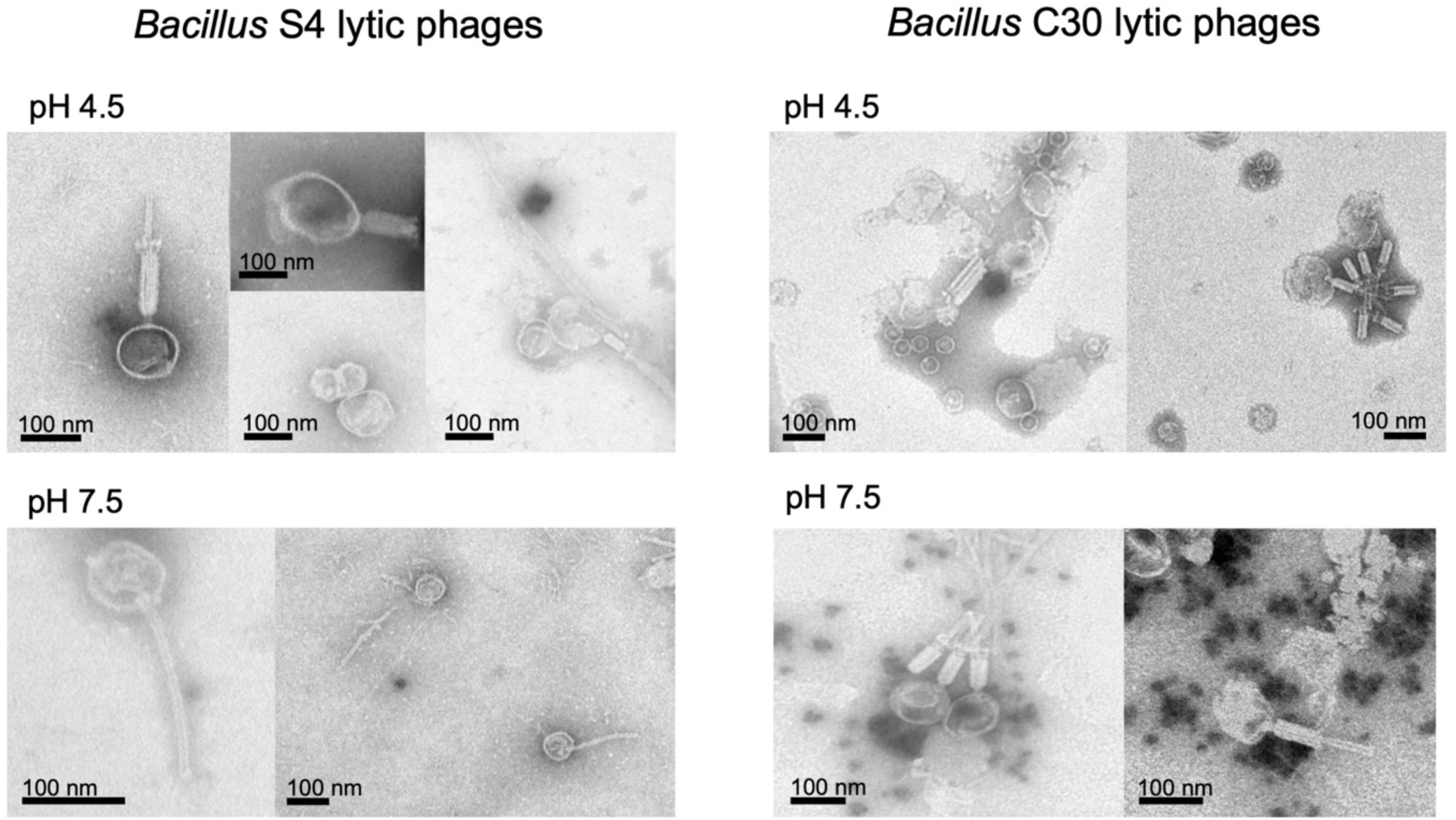
Transmission electron microscopy of phage populations from pH 4.5 and 7.5 soil samples obtained from plaque assays with *Bacillus* S4 and C30 hosts.

